# Novel Type 2-high gene clusters associated with corticosteroid sensitivity in COPD

**DOI:** 10.1101/2020.01.22.912022

**Authors:** A. Faiz, S. Pavlidis, C.S. Kuo, A. Rowe, P. S. Hiemstra, W. Timens, M. Berg, M. Wisman, Y. Guo, R. Djukanovic, P. J. Sterk, M.C. Nawijn, I. M. Adcock, K. F. Chung, M. van den Berge

## Abstract

**Rationale:** Severe asthma and COPD share common pathophysiologic traits such as relative corticosteroid insensitivity. We recently published three transcriptome-associated clusters (TACs) using hierarchical analysis of the sputum transcriptome in asthmatics from the U-BIOPRED cohort with one Type 2-high signature (TAC1) and 2 Type 2-low signatures.

**Objective:** We examined whether gene-expression signatures obtained in asthma can be used to identify the subgroup of COPD patients with steroid sensitivity.

**Methods:** Using gene set variation analysis (GSVA), we examined the distribution and enrichment scores (ES) of the 3 TACs in the transcriptome of bronchial biopsies from 46 patients who participated in the GLUCOLD COPD study that received 30 months of treatment with inhaled corticosteroids (ICS) with and without an added long-acting β-agonist (LABA). The identified signatures were then associated with longitudinal clinical variables after treatment.

**Measurements and main results:** Bronchial biopsies in COPD patients at baseline showed a wide range of expression of the 3 TACs. After ICS±LABA treatment, the ES of TAC1 was significantly reduced at 30 months, but those of TAC2 and TAC3 were unaffected. A corticosteroid-sensitive TAC1 (sub)signature was developed from the TAC1 ICS-responsive genes. This signature consisted of mast cell-specific genes identified by single-cell RNA-seq and positively correlated with bronchial biopsy mast cell numbers following ICS±LABA. Baseline levels of gene transcription predicted change in FEV1 %predicted following 30-month ICS±LABA.

**Conclusions:** Sputum-derived transcriptomic signatures from an asthma cohort can be recapitulated in bronchial biopsies of COPD patients and identified airway wall mast cells as a predictor of those COPD patients who will benefit from inhaled corticosteroids.

## Introduction

Severe asthma and COPD share common pathophysiologic traits such as airflow obstruction, eosinophilic inflammation and corticosteroid insensitivity (1–3). Although inhaled corticosteroids (ICS) are the gold standard therapy for controlling inflammation in asthma, patients with COPD are less likely to respond to ICS therapy (4–6). However, improvements in lung function, exacerbation risk and all-cause mortality with ICS treatment when used in combination with long-acting beta-agonist (LABA) bronchodilators have been reported recently (7, 8). In addition, some patients with COPD with no previous history of asthma can benefit from ICS, such as those with high blood eosinophil counts (9–14). It has previously been shown that gene expression profiling in bronchial biopsies in COPD may identify those with asthma-COPD overlap, as reflected by a Th2-high inflammatory profile and a favorable short- and long-term ICS treatment response (3).

We have recently described three transcriptome-associated clusters (TACs) in the U-BIOPRED asthma cohort using hierarchical clustering of genes differentially expressed in sputum between eosinophil-high and eosinophil-low asthmatics (15). These TACs consisted of an interleukin (IL)-13/T2-high predominantly eosinophilic cluster (TAC1) and two T2-low gene expression clusters characterized by neutrophilic inflammation and inflammasome activation (TAC2), and metabolic and mitochondrial pathways (e.g. Mitochondrial Oxidative Phosphorylation (OXPHOS)) (TAC3). These TAC signatures define patients with distinct clinical phenotypes. TAC1 was associated with sputum eosinophilia as well as elevated exhaled nitric oxide levels and was restricted to patients with severe asthma, frequent exacerbations and severe airflow obstruction. Given that these patients were dependent on corticosteroids, we hypothesize that TAC1 identifies a steroid-sensitive subphenotype of severe asthma. TAC2 showed the highest sputum neutrophilia, serum C-reactive protein levels and prevalence of eczema, while TAC3 asthma patients had normal to moderately elevated sputum eosinophils and better lung function (15).

In the current study, we hypothesized that these TAC signatures are also expressed in bronchial biopsies of mild-to-moderate COPD patients. Initially, we determined whether we could remap the sputum derived TAC signatures to biopsy transcriptional data. To do this we performed an unsupervised clustering of the genes, previously associated with the sputum TAC signatures, in the bronchial biopsy transcriptome. Using gene clustering, we derived the same TAC signatures in COPD patients. Next, we investigated the influence of short-term (0-6 months) and long-term (0-30 months) ICS therapy on these signatures as well as their ability to predict corticosteroid responsiveness in COPD.

## Materials and Methods

### Patients and study design

#### GLUCOLD Study

Gene-expression profiling was performed in bronchial biopsies from COPD patients participating in the Groningen Leiden Universities Corticosteroids in Obstructive Lung Disease (GLUCOLD) study (16). The study was approved by the local medical ethics committee and all patients provided their written informed consent. The study design and inclusion criteria have been previously described (16). Briefly, this study required participants to be either current or ex-smokers with COPD at GOLD stages 2 and 3 and not to have used ICS treatment for at least 6 months prior to entry to the study. There were 4 parallel groups of patients who were treated with 1) placebo twice daily for 30 months (Placebo group), or 2) Fluticasone propionate (FP) 500µg twice daily for 30 months (FP group), or 3) FP 500µg + Salmeterol (S) 50µg twice daily for 30 months (FP/S group), or 4) FP 500µg twice daily for 6 months and then 24 months with placebo (FP/Placebo group), referred to as the withdrawal group. As we have previously shown there is a minimal effect of the LABA in the presence of ICS on bronchial biopsy gene expression compared to ICS alone (9), groups 2 and 3 were analyzed together and will be referred to as ICS±LABA group. Bronchial biopsies were taken at baseline and after 6- and 30-months, amongst others for microarray gene-expression profiling. The methods for mRNA isolation, labeling, microarray hybridization (Affymetrix HuGene ST1.0 arrays) and data processing have been described previously (9, 17).

#### Sputum collection

Sputum was induced by inhalation of hypertonic saline solution and sputum plugs were collected from which sputum cells and sputum supernatants were obtained, as described previously [16]. Expression profiling was performed using Affymetrix U133 Plus 2.0 (Affymetrix, Santa Clara, CA, USA) microarrays with RNA extracted from sputum cells, as described previously (15).

#### Signatures summarized by gene set variation analysis

Gene Set Variation Analysis (GSVA) was used to calculate composite scores of the previously described sputum TAC signatures (table S1)(18) within the GLUCOLD bronchial biopsy transcriptome data as indicated by a sample-wise enrichment score (ES). ANOVA was used to analyse the ES differences among group means and a t-test was applied to compare the ES differences between the two means.

#### Clustering

To recapitulate the TAC signatures in the bronchial biopsies of the GLUCOLD study, hierarchical clustering analysis was conducted on the genes previously associated with the sputum TAC1, TAC2 and TAC3 signatures in bronchial biopsies at baseline, using the method as previously described (15). The intensities of the raw probe sets were log2 transformed and normalised by the robust multiarray average method (19).

#### Correlation of TAC signatures with inflammatory and clinical parameters

Baseline TAC signatures in bronchial biopsies (TACs) were correlated with each other and baseline sputum and biopsy inflammatory cell counts as well as treatment-induced changes in inflammatory cell counts and lung function measurements using Spearman correlation.

#### Bronchial biopsy single-cell signatures

Single-cell gene expression signatures for 14 clusters representing cell types or subsets (Activated endothelium, B cells, Basal 1, Basal 2, Basal activated, Basal cycling, Ciliated, Club, Cycling, DCs, Fibroblasts, Goblet, Inflammatory DCs, Ionocytes, Luminal macrophages, Mucous ciliated, Neutrophils, Resting endothelium, Smooth muscle, Serous cells from submucosal glands, T cells and Mast cells) were taken from our previously-published data to determine shifts in cell type composition using gene expression levels (20). Cell-type specific expression signatures were created by determining the percentage of reads each cell type contributes to the overall counts for a given gene. Genes with counts<200 across all cell types were removed from the analysis. Genes with >50% of total reads mapping to a single cell type and <10% reads in any other cell type were selected as cell-type-specific genes to generate the signatures. A maximum of 25 genes was taken for single types that had >25 unique genes. As not all genes detected in the single-cell seq were present on the Affymetrix hu gene ST1.0 microarray, genes not present on the array were removed from each signature. A heatmap was made of the genes corresponding to the identified cell types using gene expression dataset from the GLUCOLD study. To investigate the influence of ICS on mast cells an ES score was created for each cell type using GSVA.

#### Air Liquid Interface culture (ALI) and RNA-Seq

Primary airway epithelial cells (n=6) were cultured as previously described (21, 22). Following differentiation, for 28 days cells were then growth factor deprived overnight and then treated apically with/without Fluticasone Propionate (10^-8^M) for 24 hours. RNA was then extracted and processed for RNA-Seq as previously described (22). Differential gene expression analysis was done using the R package limma (V3.42.0) correcting age gender and the first 2 principal components.

## Results

### Patient Demographics

A total of 81 out of 89 randomized COPD patients from the placebo (n=23), ICS±LABA 30-month treatment groups (n=39) and ICS withdrawal group (n=21) had bronchial biopsy RNA available of sufficient quality to be run on the Affymetrix Hugene ST1.0 arrays. Table S2 shows the patient demographics.

### Clustering of the TAC signatures in bronchial- and sputum-derived TAC phenotypes

Initially, we performed hierarchical clustering on the genes associated with all three TAC signatures in bronchial biopsies at baseline (n=58) (**Figure 1A**); the withdrawal group (group 4) was removed for this analysis to avoid batch effects as these arrays were run at a different time point. Three distinct subgroups of COPD patients were identified (clusters green, blue and red, corresponding to TAC1, TAC2 and TAC3). We next used GSVA to create enrichment scores for several biological pathways, which previously distinguished each original sputum derived TAC signature (15). We compared the enrichment of these pathways in the three biopsy TACs (bTACs) of the COPD patients. The red cluster, bTAC1, was associated with an IL13/T2 signature (**Figure 1B**), which was previously found to distinguish the eosinophil high sputum TAC1 signature (15). The green cluster, bTAC2, was associated with the inflammasome signature (**Figure 1C**), which was previously found to distinguish the neutrophil high sTAC2 signature. No individual group was found to be solely associated with oxidative phosphorylation (OXPHOS) (**Figure 1D**), a pathway that was characteristic of sTAC3 in asthma. As all COPD patients were either current- or ex-smokers, we also investigated a smoking-related signature, derived from bronchial brushes of current and non-smokers (23). This smoking-related signature was found to distinguish bTAC3 in out COPD cohort from the sTAC3 in the asthma study (**Figure 1E**), and to be associated with the COPD bronchial-derived blue cluster (**Figure 1F**). Here we show that the same TAC signatures derived from asthma sputum can be recapitulated in COPD bronchial biopsies. Henceforth, we will use the signatures derived from the TAC publication (15).

**Figure 1.**
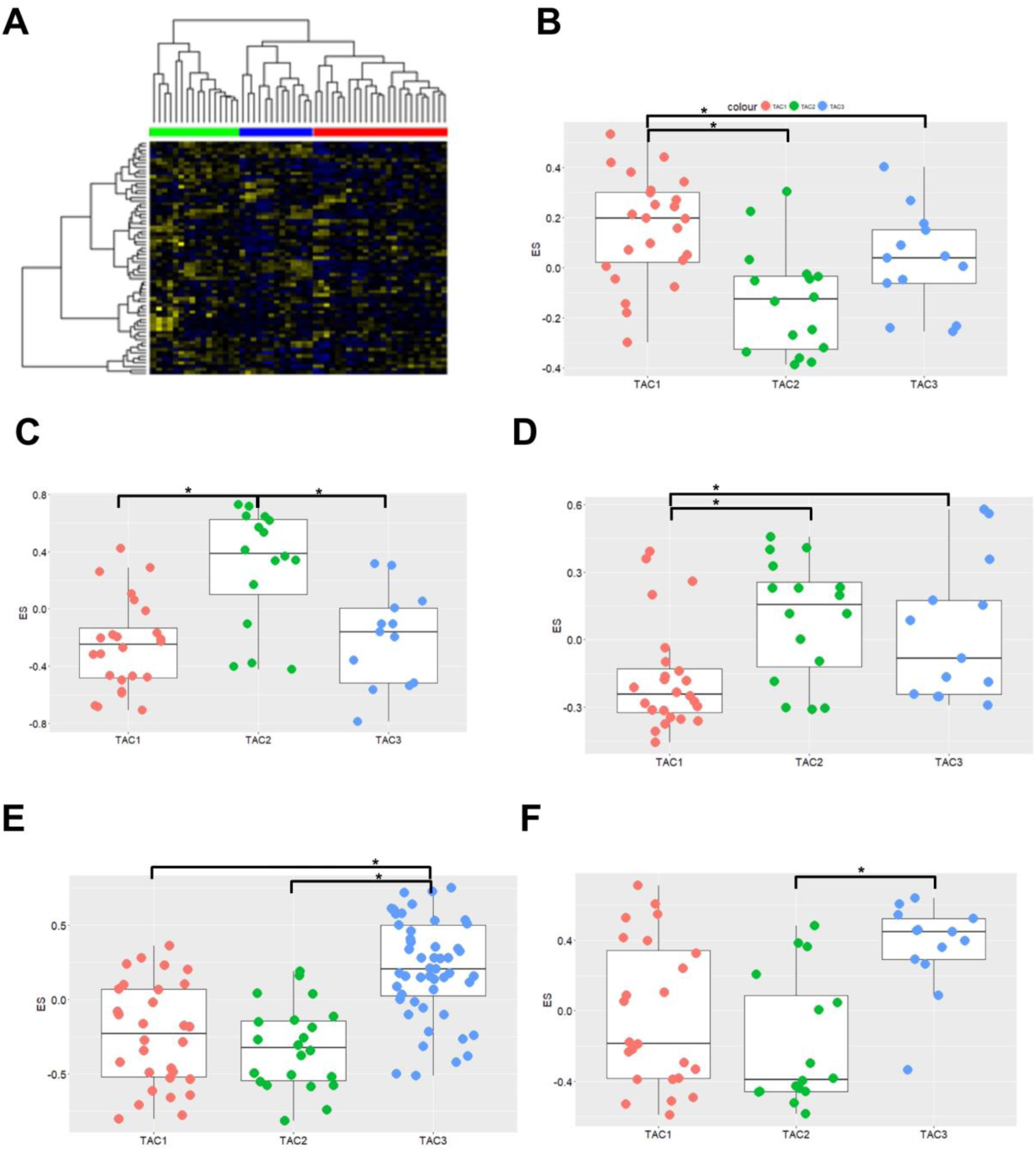
Identification of transcriptome-associated cluster (TAC) signatures in bronchial biopsies. A) Heatmap of genes that best discriminate each bronchial-derived TAC (TAC) signature (n=58). Columns represent COPD subjects and rows represent genes. Association of TAC groups 1-3 with gene signatures derived from IL13/TH2-stimulated epithelial cells (Panel B), Inflammasome activation (Panel C), oxidative phosphorylation (OXPHOS) (Panel D), cigarette smoke up-regulated signature in sputum samples from UBIOPRED (Panel E) and cigarette smoke irreversibly up-regulated signatures in bronchial brushes in GLUCOLD TAC groups (Panel F). Students unpaired T-test was used to compare each TAC group. *=p<0.05 Abbreviations: ES= enrichment score

### Relationship between TAC signatures

ES scores were created for all three TACs using the bronchial biopsy expression dataset at baseline. Bronchial biopsies at baseline showed a wide range of expression of the 3 TACs (**Figure 2A**), with a significant inverse correlation between ES TAC1 and ES TAC3 (r=-0.3814, N=58, p=0.0031) and a positive correlation between ES TAC2 and ES TAC3 (r=0.4789, N=58, p=0.0001, **Figure 2B-D**). No significant correlation between ES TAC1 and ES TAC2 was observed. These findings indicate that there is a relationship between the TAC signatures and they are not independent features.

**Figure 2.**
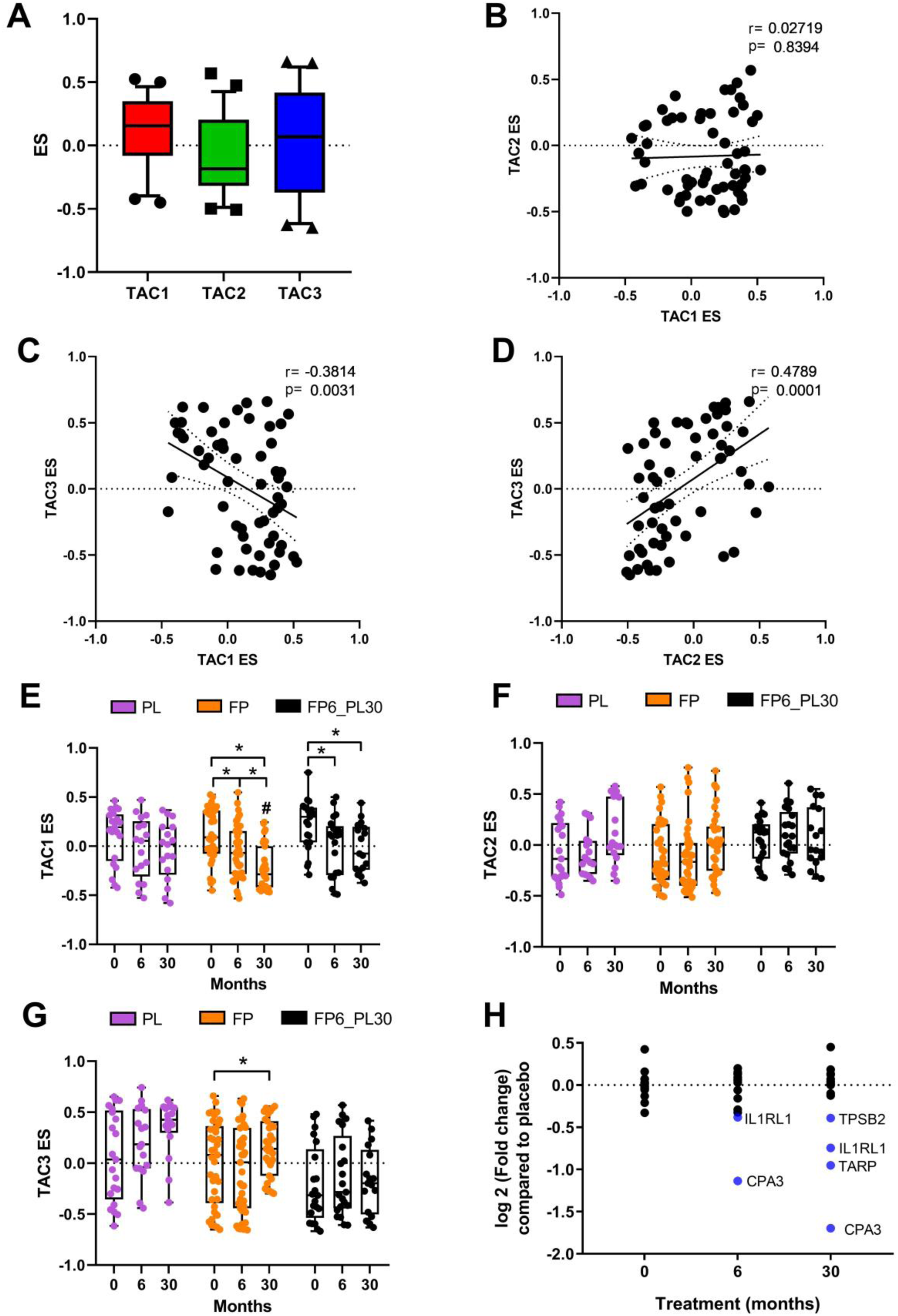
Relationship between bronchial-derived transcriptome-associated cluster (TAC) signatures and the influence of inhaled corticosteroid (ICS) treatment. A) Boxplot of the expression of TAC signatures across all patients at baseline (median±95% CI). Correlations of the expression signatures between B) TAC1 and TAC2, C) TAC1 and TAC3 and D) TAC2 and TAC3 (n=58). The influence of ICS treatment on E) TAC1, B) TAC2 and F) TAC3. D) The influence of ICS treatment on ICS sensitive TAC1 signature. G) Fold change of TAC1 genes comparing bronchial expression profiles of placebo and treatment arms of the GLUCOLD study at baseline, 6 months and 30 months. Blue represents significant down-regulated genes (adjusted pvalue<0.05). *=p<0.05 Abbreviations: PL= Placebo, FP= Fluticasone propionate ± Salmeterol and FP6_PL30= FP for 6 months and then 24 months with placebo. Abbreviations: ES= enrichment score

### Longitudinal treatment with ICS modulates the TAC1 signature only

We next investigated whether the ES of the TAC signatures were altered during ICS treatment and after ICS withdrawal (**Figure 2E**). The ICS withdrawal arm of the study was only compared to itself due to batch effects in the arrays used to quantify gene expression levels. The ES of the TAC1 signature was found to be reduced following 6 and 30 months of ICS±LABA treatment compared to baseline. No differences were seen in the placebo group over 30 months. However, following 24 months of ICS withdrawal, the ES value did not return to baseline to baseline. ICS treatment had no effect on the inflammasome-TAC2 ES but did increase TAC3 in the 30 months ICS group, but this was not significant compared to the time-matched placebo group (**Figure 2F-G**). A differential gene expression analysis (**Figure 2H**, Table S3) indicated that the ICS-induced shift in the TAC1 signature was driven by four genes: Interleukin 1 Receptor Like 1 (IL1RL1), TCR Gamma Alternate Reading Frame Protein (TARP), Tryptase beta-2 (TPSB2) and carboxypeptidase A3 (CPA3). To determine which cell-types were contributing to TAC1 sensitivity to ICS we applied our previously published single-cell RNA sequencing (scRNA-Seq) dataset obtained from bronchial biopsies of healthy control and asthmatic patients (20). Initially, tSNE plots were created for the ICS sensitive genes (IL1RL1, TPSB2 and CPA3) from

TAC1, while TARP was not present in the scRNA-Seq dataset. tSNE plots made from epithelial cell subpopulations/subsets showed no specific expression of the three genes (**Figure S1A-D**), while the tSNE plot of non-epithelial cell populations identified all three genes as mast cell-specific (**Figure 3A-D**). To further investigate the effect of ICS on mast cells and other cell populations, we took two approaches: 1) we examined the change in gene expression of cell type-specific markers developed from the scRNA-seq data obtained from bronchial biopsies, and 2) we examined the histological staining of mast cells in adjacent biopsies from the same patients (20). Of the 14 cell type-specific gene signatures, the mast cell signature was found to be one of the most sensitive to ICS treatment (**Figure 3E**). A GSVA analysis of mast cell-specific genes identified from the single-cell seq (TPSB2, SLC18A2, MS4A2, HPGDS, and CPA3) shows a clear decrease in the mast cell signature after 6 and 30 months ICS±LABA treatment compared to placebo, with an increase in expression of this signature following 24 month ICS withdrawal (**Figure 3F**). Similar results were found for the histological staining of mast cells cell numbers measured by tryptase (AA1) staining in adjacent biopsies (**Figure 3G**). Moreover, a decrease was seen in the placebo arm at 6 and 30 months but this was not significantly lower compared to time-matched ICS treatment controls. There was a statistically significant correlation between the mast cell signature and log mast cell counts in adjacent biopsies (**Figure S2**). Finally, we investigated the percentage that each cell type contributes to the overall gene expression of CPA3, TPSB2 and IL1RL1 in bronchial biopsies (**Figure 4A**). Although IL1RL1 does appear to be mast cell-specific in tSNE plots, more than ∼50% of its total expression in biopsies is coming from basal epithelial cells. To determine whether the decrease of IL1RL1, TPSB2 and CPA3 was a direct effect of corticosteroid therapy, we treated airway epithelial cells differentiated at air-liquid interface with Fluticasone Propionate (FP) for 24 hours and measured gene expression by RNA-Seq. IL1RL1 gene expression was found to be significantly decreased by FP, while CPA3 and TPSB2 were not detected in airway epithelial cells, which is to be expected as both CPA3 and TPSB2 were specifically expressed in the mast cell cluster base on single-cell Seq (**Figure 4B**).

**Figure 3.**
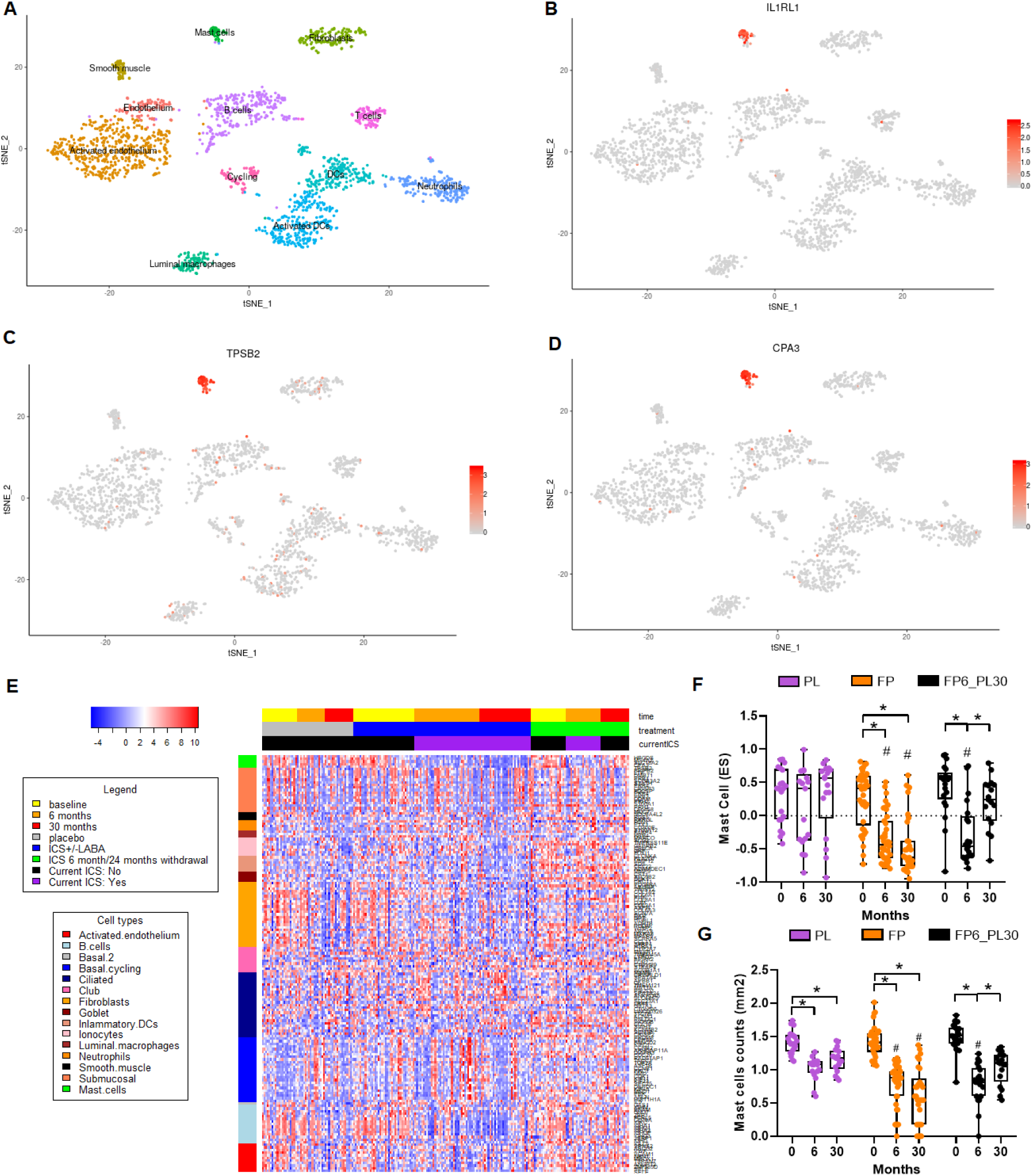
Determining the basis of ICS sensitivity of the TAC1 signature. TSNE plots for ICS sensitive TAC1 genes (IL1RL1, TPSB2 and CPA3), obtained from of single cell seq data obtained from asthmatic (n=4) and healthy controls (n=4) (A-D). E) Heatmap of cell population-specific genes identified by single-cell sequencing from bronchial biopsies. F) GSVA analysis of the mast cell signature derived from single cell-sequencing. G) Plot of log mast cell counts in bronchial biopsies (mm2) treated with either Placebo, ICS for 6 month then 24-month withdrawal or ICS±LABA for 30 months. Abbreviations: PL= Placebo, FP= Fluticasone propionate ± Salmeterol and FP6_PL30= FP for 6 months and then 24 months with placebo. *=p<0.05 and #=p<0.05 compared to time-matched placebo treatment using linear mix effects models using patient ID as the random variable. Abbreviations: ES= enrichment score

**Figure 4.**
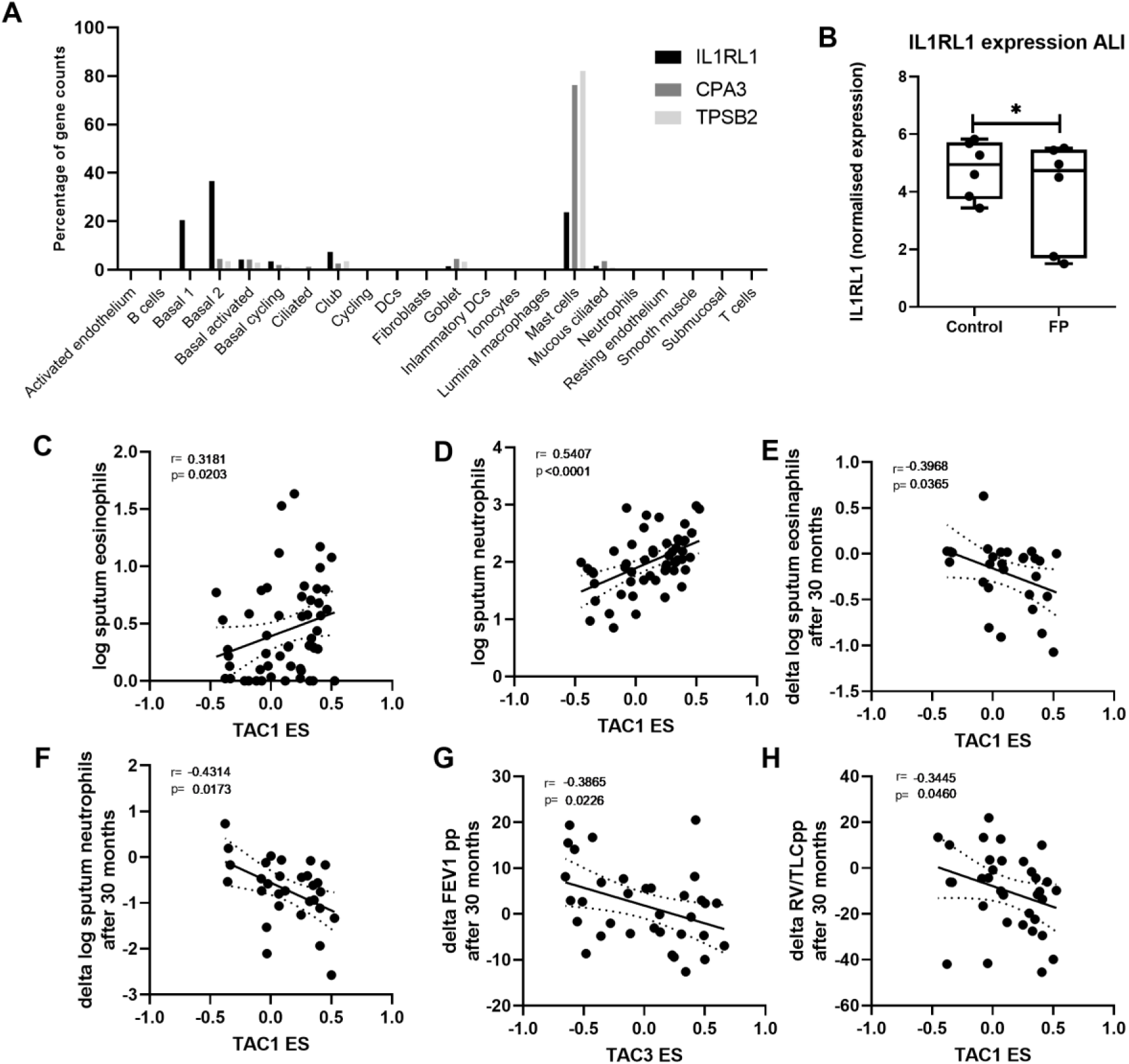
Correlation of bronchial-derived transcriptome-associated cluster (TAC) signatures and inflammatory cell counts. F) IL1RL1 expression from primary airway epithelial cells grown at air-liquid interface, quiecesed overnight and then treated with Fluticasone propionate (FP; 10^-8^ M) for 24 hours (N=6 donors). Correlation of TAC1 signature at baseline with sputum A) log eosinophil counts and B) log neutrophil counts. Correlation of TAC1 signature at baseline with C) change in log eosinophil counts and D) delta log neutrophil counts after 30 months ICS ±LABA. E) Correlation of TAC3 signature at baseline with delta FEV1% predicted F) Correlation of ICS-sensitive TAC1 signature at baseline with delta FEV1 % predicted.

### TAC association with physiologic and inflammatory features of COPD

To determine the relationship between the TAC signatures and clinical and inflammatory features, we first investigated their relationship to matched sputum inflammatory cell profiles. At baseline, higher TAC1 was associated with higher sputum eosinophil and neutrophil counts (**Figure 4C&D**). The ES of TAC1-3 was not associated with airflow obstruction (FEV1% predicted) or hyperinflation (residual volume, RV %predicted) (**Table 1**). Finally, we determined whether baseline TAC signatures could predict the ICS treatment response as reflected by improvement in lung function and decrease in inflammatory cell counts. Higher TAC1 baseline ES was associated with a more pronounced decrease in sputum neutrophil and eosinophil numbers after 30 months treatment with ICS±LABA (**Figure 4E&F**, **Table 2**), while a higher TAC3 was associated with lesser improvement of lung function measured by FEV1% predicted (**Figure 4G**). Interestingly, TAC1 signature at baseline, was associated with improvement of lung function measured by RV/TLC% predicted over the 30-month period (**Figure 4H**).

**Table 1.** Association of TAC signatures with clinical variables at baseline

|  | TAC1 |  | TAC2 |  | TAC3 |  |
| --- | --- | --- | --- | --- | --- | --- |
|  | rho | P value | rho | P value | rho | P value |
| Log sputum eosinophil counts | **0.3181** | **0.0203** | -0.1503 | 0.2828 | -0.2281 | 0.1005 |
| Log sputum neutrophil counts | **0.5407** | **<0.0001** | 0.02508 | 0.8585 | **-0.2977** | **0.0304** |
| Log sputum macrophage counts | **0.3391** | **0.013** | -0.0125 | 0.9292 | -0.09136 | 0.5153 |
| Log sputum lymphocyte counts | **0.502** | **0.0001** | 0.08958 | 0.5235 | -0.2358 | 0.0891 |
| FEV<sub>1</sub> %predicted | -0.0935 | 0.4852 | 0.01461 | 0.9133 | 0.1402 | 0.2939 |
| RV/TLC %predicted | 0.0064 | 0.9618 | -0.0707 | 0.5978 | -0.08069 | 0.5471 |

Abbreviations: FEV<sub>1</sub> = Forced Expiratory Volume in 1second and RV/TLC= residual volume/total lung capacity

**Table 2.** Predictive value of baseline TAC signatures to predict change of clinical variables over 30 months ICS+/-LABA treatment compared to baseline.

|  | TAC1 |  | TAC2 |  | TAC3 |  |
| --- | --- | --- | --- | --- | --- | --- |
|  | rho | P value | rho | P value | rho | P value |
| FEV<sub>1</sub> %predicted | -0.0414 | 0.8161 | -0.0778 | 0.662 | **-0.3901** | **0.0226** |
| RV/TLC %predicted | **-0.3445** | **0.046** | 0.2131 | 0.2262 | 0.2648 | 0.1302 |
| Log sputum eosinophils counts | **-0.3968** | **0.0365** | 0.1642 | 0.4037 | 0.3454 | 0.0718 |
| Log sputum neutrophil counts | **-0.4314** | **0.0173** | 0.0309 | 0.8711 | 0.2405 | 0.2005 |

Abbreviations: FEV<sub>1</sub> = Forced Expiratory Volume in 1second and RV/TLC= residual volume/total lung capacity

## Discussion

We found that the genes represented in the TAC signatures derived from the sputum of the U-BIOPRED asthma cohort are also expressed in bronchial biopsies of COPD patients. High enrichment of the TAC1 signature (high eosinophilia/Th2 high) in COPD serves to confirm the existence of an eosinophilic phenotype associated with the T2 pathway. In addition, the suppression of this signature by inhaled corticosteroid therapy and the reversal of the expression of this signature by cessation of inhaled corticosteroid therapy reinforces the similarity of this COPD endotype to severe eosinophilic asthma. Furthermore, we found that the ICS treatment effect on TAC1 was the consequence of three genes expressed in mast cells: TPSB2 (a tryptase II gene, selectively expressed in mast cells), IL1RL1 (the receptor for IL-33) which is expressed on many inflammatory cell types including mast cells, eosinophils, neutrophils, innate lymphoid cells and epithelial cells and CPA3 (carboxypeptidase A3), a gene selectively expressed in mast cells, and part of a sputum signature that predicts treatment response in asthma (24, 25). Finally, the TAC1 signature correlated with sputum eosinophil and neutrophil counts and also predicted ICS response at both 6 and 30 months in COPD. In contrast, gene signatures associated with the TAC2 (inflammasome) and TAC3 (smoking signature) were unaffected by ICS therapy or its subsequent withdrawal.

An increased TAC1 expression, previously identified to reflect Th2 high inflammation, was found to be associated with higher sputum neutrophil and eosinophil counts. These latter results are in line with previous findings that asthma-derived Th2 signatures measured in bronchial biopsies correlate with airway wall eosinophil counts and blood eosinophil percentages and a more severe airflow obstruction in COPD (3). The association with TAC1 and neutrophils has not been described in previous publications and may be due to the difference in gene expression levels and cell composition between the original sputum samples and the biopsy samples. Furthermore, we show that higher baseline TAC1 expression predicts a more pronounced ICS treatment-induced decrease in eosinophil and neutrophil numbers in sputum. Furthermore, TAC1 predicted an ICS-induced improvement in RV/TLC% predicted. The ICS sensitivity of the original TAC1 signature was predominantly associated with the genes CPA3 and IL1RL1. CPA3 is a metalloprotease usually used as a activation marker of mast cells (26), while IL1RL1 has been identified in a number of Genome-Wide Association Studies (GWAS) to be associated with wheezing phenotypes and asthma in childhood (27), with the risk allele being associated with higher expression of the gene in airway tissue (28). Both genes are associated with the Th2 response in asthma (18).

Interestingly in asthma, it has previously been shown that the severity of the diseases as measure by hyperresponsiveness in the presence of corticosteroid treatment was related to Mast cell % in bronchial brushes (29). While in COPD, Tryptase positive mast cells were found to be positively correlated with lung function (FEV1/VC) (30). In the current study, we show that the TAC1 signature which is driven by mast cell-specific genes at baseline can predict improvement in RV/TLC% predicted following 30 months ICS therapy.

Mast cells reside in the airways and other organs and impact on both the innate and adaptive immune response via the secretion of numerous inflammatory bronchoconstrictor mediators such as leukotrienes and prostaglandins (31). Activated mast cells, particularly chymase and tryptase positive connective tissue mast cells, release large amounts of CPA3. Interestingly, IL1RL1 is not only present on granulocytes but also expressed constitutively by mast cells and its ligand, IL-33, stimulates mast cell adhesion to laminin, fibronectin, and vitronectin, and mast cell survival, growth, development, and maturation (32). In addition, mast cell-derived tryptase and chymase cleave IL-33 to generate mature active forms (32). Together, these data suggest that mast cells represent a major target for ICS in ICS-sensitive COPD patients. However, whether this shift in mast cells is a consequence of the causal mechanism, not the mechanism itself remains to be determined. Although we did find IL1RL1 expression to decrease by corticosteroids directly in the airway epithelium, the lower expression in bronchial biopsies may result from the decrease in mast cell numbers and only partially be due to direct repression of gene expression in other cell types. Furthermore, we are blinded to the eosinophils that are also known to express IL1RL1 due to the difficulty of performing sc-Seq on this cell type (20).

The TAC signatures, originally derived from the U-BIOPRED asthma cohort, were recapitulated in bronchial biopsies and predicted corticosteroid responsiveness in patients with mild-to-moderatee COPD. This highlights a potential overlap in the pro-inflammatory signatures that are present in asthma and COPD. Asthma-COPD overlap syndrome is derived from a concept that both asthma and COPD patients can share clinical traits and therefore share responsiveness to particular treatments (1). Using asthma-derived signatures such as the TACs may provide an important method of identifying COPD patients that have asthma-like features such as responsiveness to ICS therapy, T2- or mast-cell-directed treatments.

In conclusion, we found that the sputum-derived TAC signatures could be recapitulated in bronchial biopsies of COPD patients. Furthermore, Th2-high TAC1 signature was found to be related to more sensitivity to ICS and to be predictive for the degree of suppression of mast cell inflammation following ICS treatment. Finally, this study shows that asthma and COPD share common gene signatures, which may be used to predict subsets of COPD patients sensitive to corticosteroids.

## Acknowledgments

The U-BIOPRED consortium received funding from the European Union and from the European Federation of Pharmaceutical Industries and Associations as an Innovative Medicines Initiative Joint Undertaking funded project (115010) on behalf of the U-BIOPRED Study Group with input from the U-BIOPRED Patient Input Platform and patient representatives from the Ethics Board and Safety Management Board. SP was supported by the EU-EFPIA Innovative Medicines Initiative eTRIKS project (115446). KFC is a Senior Investigator of the UK National Institute for Health Research. IMA and KFC are PIs within the Asthma UK Centre for asthma and allergic mechanisms. This project is co-financed by the Ministry of Economic Affairs and Climate Policy by means of the PPP Allowance made available by the Top Sector Life Sciences & Health to stimulate public-private partnerships. The GLUCOLD study was was funded by unrestricted grants from the Stichting Astma Bestrijding, the Netherlands Asthma Foundation, Netherlands Organization for Scientific Research (ZonMw), GlaxoSmithKline, the Royal Dutch Academy of Sciencesand Arts, the University Medical Center Groningen and Leiden University Medical Center and the NIH R01 HL095388 (Spira/Lenburg).

## Supplementary

**Figure S1.**
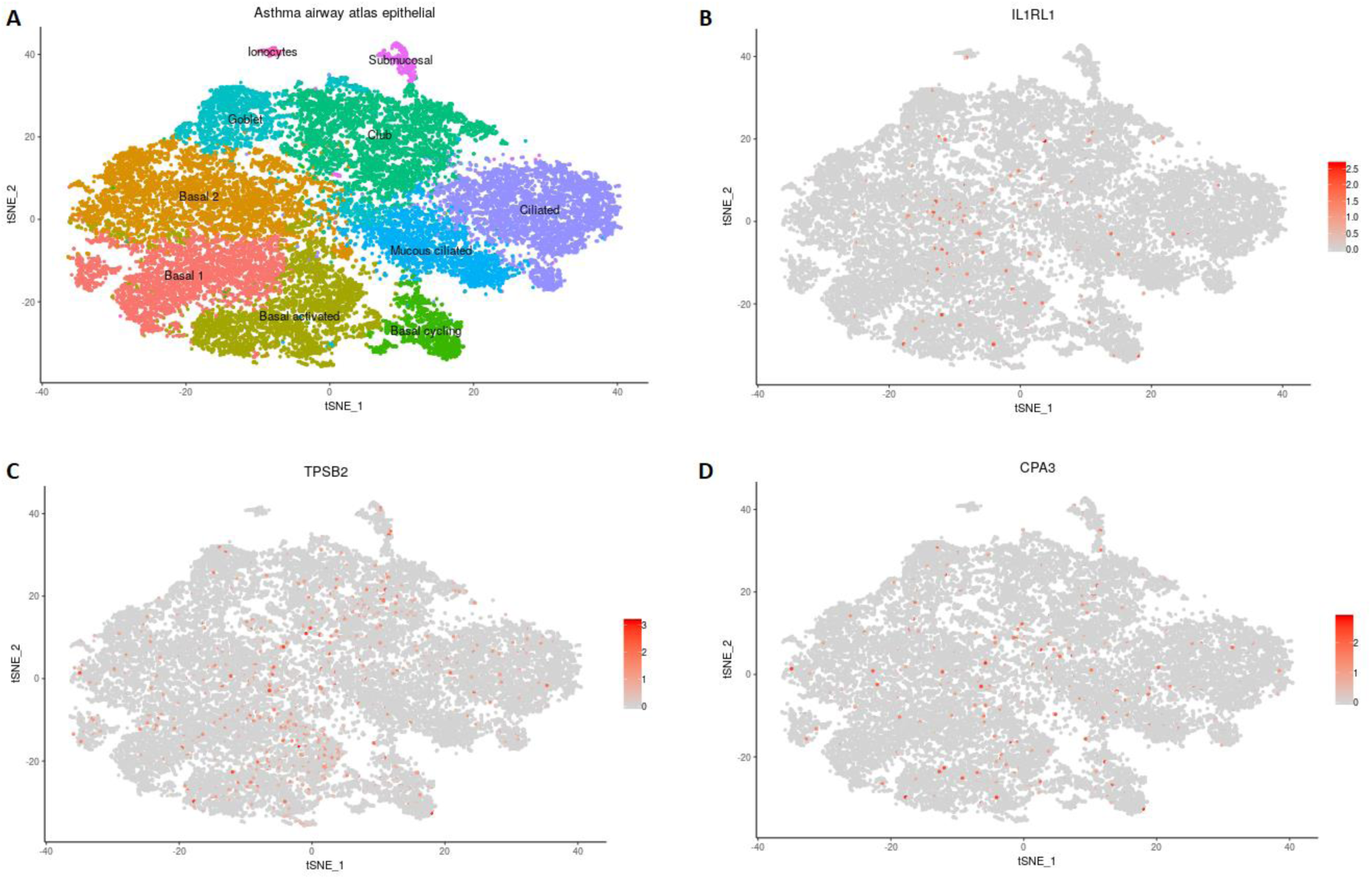
TSNE plots made from epithelial only cell/subtypes. TSNE plots for ICS sensitive TAC1 genes (IL1RL1, TPSB2 and CPA3), obtained from of single cell seq data obtained from asthmatic (n=4) and healthy controls (n=4) (A-D).

**Table S1.**
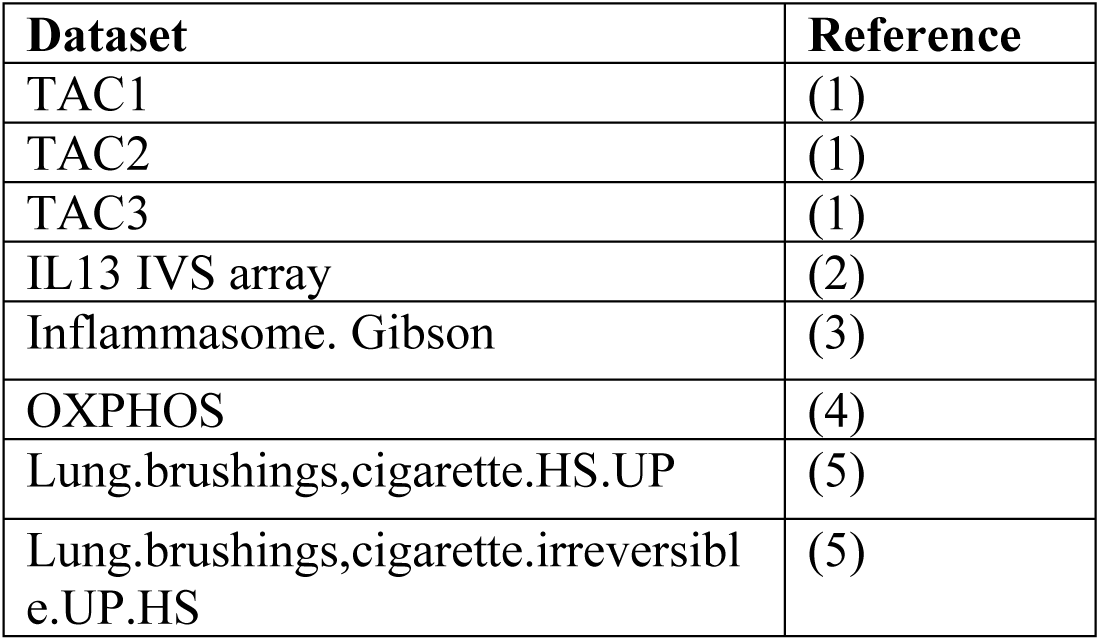
Gene set variation analysis (GSVA) datasets.

**Table S2.** GLUCOLD patient characteristics.

|  | **Fluticasone±salmeterol for 30 months** |  |  | **Placebo for 30 months** |  |  | **Fluticasone for 6 months followed by placebo between 6 and 30 months** |  |  |
| --- | --- | --- | --- | --- | --- | --- | --- | --- | --- |
|  | **Baseline** | **6 months** | **30 months** | **Baseline** | **6 months** | **30 months** | **Baseline** | **6 months** | **30 months** |
| Number of included patients | 45 |  |  | 23 |  |  | 21 |  |  |
| Number of biopsies available at each time point | 37 | 39 | 31 | 21 | 17 | 17 | 21 | 21 | 17 |
| Male/female, n | 41/4 |  |  | 19/2 |  |  | 19/2 |  |  |
| Age, years | 62.4±7.2 |  |  | 60.2±7.8 |  |  | 63.1±7.4 |  |  |
| BMI | 25.5±3.7 |  |  | 24.2±3.9 |  |  | 25.4±3.6 |  |  |
| Current smokers, n (%) | 22 (59) | 20 (51) | 14 (45) | 14 (67) | 10 (59) | 8 (47) | 10 (48) | 9 (43) | 8 (47) |
| RIN score | 3.3±1.5 | 3.5±1.3 | 4.8±1.5** | 3.5±1.3 | 3.9±1.6 | 5.2±1.8** | 3.3±1.7 | 3.7±1.7 | 3.7±1.5 |
| FEV<sub>1</sub>, %predicted | 62.6±9.0 | 63.6±10.7 | 64.2±12.3 | 61.3±8.80 | 62.3±9.20 | 57.0±8.3 | 64.7±8.62 | 64.9±9.0 | 64.2±12.5 |

**Table S3.** TAC1 differential gene expression analysis comparing bronchial expression profiles of placebo and treatment arms of the GLUCOLD study at baseline, 6 months, and 30 months.

|  | Baseline |  |  | 6 months |  |  | 30 months |  |  |
| --- | --- | --- | --- | --- | --- | --- | --- | --- | --- |
|  | logFC | P Value | FDR | logFC | P Value | FDR | logFC | P Value | FDR |
| CPA3 | -0.205 | 4.159E-01 | 9.221E-01 | -1.142 | 2.871E-05 | 4.593E-04 | -1.702 | 3.421E-09 | 5.473E-08 |
| IL1RL1 | -0.036 | 7.815E-01 | 9.221E-01 | -0.384 | 5.495E-03 | 4.396E-02 | -0.749 | 3.050E-07 | 2.440E-06 |
| TARP | -0.330 | 1.322E-01 | 5.286E-01 | -0.329 | 1.558E-01 | 2.769E-01 | -0.958 | 8.398E-05 | 4.479E-04 |
| TPSB2 | 0.032 | 8.073E-01 | 9.221E-01 | -0.293 | 3.914E-02 | 1.566E-01 | -0.391 | 7.909E-03 | 3.164E-02 |
| ATP2A3 | 0.071 | 6.903E-01 | 9.221E-01 | 0.029 | 8.763E-01 | 8.763E-01 | 0.453 | 2.040E-02 | 6.529E-02 |
| LGALS12 | 0.158 | 6.693E-02 | 5.246E-01 | 0.201 | 2.871E-02 | 1.531E-01 | 0.185 | 5.122E-02 | 1.366E-01 |
| PRSS33 | -0.017 | 8.142E-01 | 9.221E-01 | 0.135 | 8.707E-02 | 2.322E-01 | 0.128 | 1.170E-01 | 2.675E-01 |
| OLIG2 | -0.015 | 8.645E-01 | 9.221E-01 | 0.071 | 4.445E-01 | 5.926E-01 | 0.134 | 1.624E-01 | 3.248E-01 |
| CCR3 | 0.071 | 3.443E-01 | 9.221E-01 | 0.136 | 8.581E-02 | 2.322E-01 | 0.070 | 3.935E-01 | 6.995E-01 |
| ALOX15 | 0.428 | 2.522E-02 | 4.035E-01 | -0.060 | 7.650E-01 | 8.160E-01 | -0.092 | 6.587E-01 | 8.782E-01 |
| FAM101B | -0.056 | 4.705E-01 | 9.221E-01 | 0.058 | 4.846E-01 | 5.965E-01 | 0.042 | 6.185E-01 | 8.782E-01 |
| CD24 | 0.016 | 9.292E-01 | 9.292E-01 | -0.156 | 4.209E-01 | 5.926E-01 | -0.117 | 5.582E-01 | 8.782E-01 |
| CLC | -0.040 | 6.655E-01 | 9.221E-01 | 0.153 | 1.158E-01 | 2.647E-01 | 0.025 | 8.065E-01 | 9.926E-01 |
| SOCS2 | -0.130 | 9.837E-02 | 5.246E-01 | 0.053 | 5.243E-01 | 5.992E-01 | -0.001 | 9.951E-01 | 9.951E-01 |
| HRH4 | 0.029 | 7.453E-01 | 9.221E-01 | 0.133 | 1.544E-01 | 2.769E-01 | 0.004 | 9.647E-01 | 9.951E-01 |
| VSTM1 | 0.012 | 8.530E-01 | 9.221E-01 | 0.093 | 1.934E-01 | 3.095E-01 | 0.008 | 9.160E-01 | 9.951E-01 |

Abbreviation FC= Fold Change, FDR=False Discovery Rate

## References

1. Postma DS, Rabe KF. The asthma–COPD overlap syndrome. New England Journal of Medicine 2015; 373: 1241–1249.

2. Cosio BG, Soriano JB, López-Campos JL, Calle-Rubio M, Soler-Cataluna JJ, de-Torres JP, Marín JM, Martínez-Gonzalez C, de Lucas P, Mir I. Defining the asthma-COPD overlap syndrome in a COPD cohort. Chest 2016; 149: 45–52.

3. Christenson SA, Steiling K, van den Berge M, Hijazi K, Hiemstra PS, Postma DS, Lenburg ME, Spira A, Woodruff PG. Asthma–COPD overlap. Clinical relevance of genomic signatures of type 2 inflammation in chronic obstructive pulmonary disease. American journal of respiratory and critical care medicine 2015; 191: 758–766.

4. Vestbo J, Søorensen T, Lange P, Brix A, Torre P, Viskum K. Long-term effect of inhaled budesonide in mild and moderate chronic obstructive pulmonary disease: a randomised controlled trial. The Lancet 1999; 353: 1819–1823.

5. Burge PS, Calverley P, Jones PW, Spencer S, Anderson JA, Maslen T. Randomised, double blind, placebo controlled study of fluticasone propionate in patients with moderate to severe chronic obstructive pulmonary disease: the ISOLDE trial. Bmj 2000; 320: 1297–1303.

6. Pauwels RA, Löfdahl C-G, Laitinen LA, Schouten JP, Postma DS, Pride NB, Ohlsson SV. Long-term treatment with inhaled budesonide in persons with mild chronic obstructive pulmonary disease who continue smoking. New England Journal of Medicine 1999; 340: 1948–1953.

7. Celli BR, Thomas NE, Anderson JA, Ferguson GT, Jenkins CR, Jones PW, Vestbo J, Knobil K, Yates JC, Calverley PMJAjor. Effect of pharmacotherapy on rate of decline of lung function in chronic obstructive pulmonary disease: results from the TORCH study. American journal of respiratory critical care medicine 2008; 178: 332–338.

8. Valipour A, Slebos D-J, Herth F, Darwiche K, Wagner M, Ficker JH, Petermann C, Hubner R-H, Stanzel F, Eberhardt R. Endobronchial valve therapy in patients with homogeneous emphysema. Results from the IMPACT study. American journal of respiratory critical care medicine 2016; 194: 1073–1082.

9. van den Berge M, Steiling K, Timens W, Hiemstra PS, Sterk PJ, Heijink IH, Liu G, Alekseyev YO, Lenburg ME, Spira A. Airway gene expression in COPD is dynamic with inhaled corticosteroid treatment and reflects biological pathways associated with disease activity. Thorax 2014; 69: 14–23.

10. Hartjes FJ, Vonk JM, Faiz A, Hiemstra PS, Lapperre TS, Kerstjens HA, Postma DS, van den Berge M, Groningen, Group LUCiOLDS. Predictive value of eosinophils and neutrophils on clinical effects of ICS in COPD. Respirology 2018; 23: 1023–1031.

11. Bafadhel M, Peterson S, De Blas MA, Calverley PM, Rennard SI, Richter K, Fagerås M. Predictors of exacerbation risk and response to budesonide in patients with chronic obstructive pulmonary disease: a post-hoc analysis of three randomised trials. The Lancet Respiratory Medicine 2018; 6: 117–126.

12. Adir Y, Hakrush O, Shteinberg M, Schneer S, Agusti A. Circulating eosinophil levels do not predict severe exacerbations in COPD: a retrospective study. ERJ open research 2018; 4: 00022–02018.

13. Roche N, Chapman KR, Vogelmeier CF, Herth FJ, Thach C, Fogel R, Olsson P, Patalano F, Banerji D, Wedzicha JA. Blood eosinophils and response to maintenance chronic obstructive pulmonary disease treatment. Data from the FLAME trial. American journal of respiratory critical care medicine 2017; 195: 1189–1197.

14. Barnes NC, Sharma R, Lettis S, Calverley PM. Blood eosinophils as a marker of response to inhaled corticosteroids in COPD. European Respiratory Journal 2016; 47: 1374–1382.

15. Kuo C-HS, Pavlidis S, Loza M, Baribaud F, Rowe A, Pandis I, Sousa A, Corfield J, Djukanovic R, Lutter R, Sterk PJ, Auffray C, Guo Y, Adcock IM, Chung KF. T-helper cell type 2 (Th2) and non-Th2 molecular phenotypes of asthma using sputum transcriptomics in U-BIOPRED. European Respiratory Journal 2017; 49: 1602135.

16. Lapperre TS, Snoeck-Stroband JB, Gosman MM, Jansen DF, van Schadewijk A, Thiadens HA, Vonk JM, Boezen HM, ten Hacken NH, Sont JK. Effect of Fluticasone With and Without Salmeterol on Pulmonary Outcomes in Chronic Obstructive Pulmonary Disease A Randomized Trial. Annals of internal medicine 2009; 151: 517–527.

17. Faiz A, Steiling K, Roffel MP, Postma DS, Spira A, Lenburg ME, Borggrewe M, Eijgenraam TR, Jonker MR, Koppelman GH. Effect of long-term corticosteroid treatment on microRNA and gene-expression profiles in COPD. European Respiratory Journal 2019; 53: 1801202.

18. Kuo C-HS, Pavlidis S, Loza M, Baribaud F, Rowe A, Pandis I, Sousa A, Corfield J, Djukanovic R, Lutter R. T-helper cell type 2 (Th2) and non-Th2 molecular phenotypes of asthma using sputum transcriptomics in U-BIOPRED. European Respiratory Journal 2017; 49: 1602135.

19. Irizarry RA, Bolstad BM, Collin F, Cope LM, Hobbs B, Speed TP. Summaries of Affymetrix GeneChip probe level data. Nucleic acids research 2003; 31: e15–e15.

20. Braga FAV, Kar G, Berg M, Carpaij OA, Polanski K, Simon LM, Brouwer S, Gomes T, Hesse L, Jiang J. A cellular census of human lungs identifies novel cell states in health and in asthma. Nature medicine 2019: 1.

21. Faiz A, Weckmann M, Tasena H, Vermeulen CJ, Van den Berge M, ten Hacken NH, Halayko AJ, Ward JP, Lee TH, Tjin GJERJ. Profiling of healthy and asthmatic airway smooth muscle cells following interleukin-1β treatment: a novel role for CCL20 in chronic mucus hypersecretion. 2018; 52: 1800310.

22. Boudewijn IM, Lan A, Faiz A, Cox CA, Brouwer S, Schokker S, Vroegop SJ, Nawijn MC, Woodruff PG, Christenson SAJA. Nasal gene expression changes with inhaled corticosteroid treatment in asthma. 2019.

23. Spira A, Beane J, Shah V, Liu G, Schembri F, Yang X, Palma J, Brody JS. Effects of cigarette smoke on the human airway epithelial cell transcriptome. Proceedings of the National Academy of Sciences 2004; 101: 10143–10148.

24. Baines KJ, Simpson JL, Wood LG, Scott RJ, Fibbens NL, Powell H, Cowan DC, Taylor DR, Cowan JO, Gibson PGJJoA, Immunology C. Sputum gene expression signature of 6 biomarkers discriminates asthma inflammatory phenotypes. 2014; 133: 997–1007.

25. Wang G, Baines KJ, Fu JJ, Wood LG, Simpson JL, McDonald VM, Cowan DC, Taylor DR, Cowan JO, Gibson PGJERJ. Sputum mast cell subtypes relate to eosinophilia and corticosteroid response in asthma. 2016; 47: 1123–1133.

26. Abonia JP, Blanchard C, Butz BB, Rainey HF, Collins MH, Stringer K, Putnam PE, Rothenberg ME. Involvement of mast cells in eosinophilic esophagitis. Journal of Allergy and Clinical Immunology 2010; 126: 140–149.

27. Savenije OE, John JMM, Granell R, Kerkhof M, Dijk FN, de Jongste JC, Smit HA, Brunekreef B, Postma DS, Van Steen K. Association of IL33–IL-1 receptor–like 1 (IL1RL1) pathway polymorphisms with wheezing phenotypes and asthma in childhood. Journal of Allergy and Clinical Immunology 2014; 134: 170–177.

28. Li X, Hastie AT, Hawkins GA, Moore WC, Ampleford EJ, Milosevic J, Li H, Busse WW, Erzurum SC, Kaminski N. eQTL of bronchial epithelial cells and bronchial alveolar lavage deciphers GWAS-identified asthma genes. Allergy 2015; 70: 1309–1318.

29. Gibson PG, Saltos N, Borgas TJJoa, immunology c. Airway mast cells and eosinophils correlate with clinical severity and airway hyperresponsiveness in corticosteroid-treated asthma. 2000; 105: 752–759.

30. Gosman MM, Postma DS, Vonk JM, Rutgers B, Lodewijk M, Smith M, Luinge MA, ten Hacken NH, Timens WJRr. Association of mast cells with lung function in chronic obstructive pulmonary disease. 2008; 9: 64.

31. Komi DEA, Grauwet K. Role of mast cells in regulation of T cell responses in experimental and clinical settings. Clinical reviews in allergy immunology 2018; 54: 432–445.

32. Chan BC, Lam CW, Tam L-S, Wong CK. IL33: Roles in Allergic Inflammation and Therapeutic Perspectives. Frontiers in immunology 2019; 10

